# TNFRSF14 (HVEM) is a novel immune checkpoint for cancer immunotherapy in humanized mice

**DOI:** 10.1101/711119

**Authors:** Nicolas Aubert, Simon Brunel, Daniel Olive, Gilles Marodon

## Abstract

**Background:** TNFRSF14 (herpes virus entry mediator (HVEM) delivers a negative signal to T cells through the B and T Lymphocyte Attenuator (BTLA) molecule and has been associated with a worse prognosis in numerous malignancies. A formal demonstration that the HVEM/BTLA axis can be targeted for cancer immunotherapy is however still lacking.

**Methods:** We used immunodeficient NOD.SCID.gc-null mice reconstituted with human PBMC and grafted with human tumor cell lines subcutaneously. Tumor growth was compared using linear and non linear regression statistical modeling. The phenotype of tumor-infiltrating leukocytes was determined by flow cytometry. Statistical testing between groups was performed by a non-parametric t test. Quantification of mRNA in the tumor was performed using NanoString pre-designed panels. Bioinformatics analyses were performed using Metascape, Gene Set Enrichment Analysis and Ingenuity Pathways Analysis with embedded statistical testing.

**Results:** We showed that a murine monoclonal antibody to human HVEM significantly impacted the growth of various HVEM-positive cancer cell lines in humanized NSG mice. Using CRISPR/cas9 mediated deletion of HVEM, we showed that HVEM expression by the tumor was necessary and sufficient to observe the therapeutic effect. Tumor cell killing by the mAb was dependent on innate immune cells still present in NSG mice, as indicated by *in vivo* and *in vitro* assays. Mechanistically, tumor control by human T cells by the mAb was dependent on CD8 T cells and was associated with an increase in the proliferation and number of tumor-infiltrating leukocytes. Accordingly, the expression of genes belonging to T cell activation pathways, such as JAK/STAT and NFKB were enriched in anti-HVEM-treated mice, whereas genes associated with immuno-suppressive pathways were decreased. Finally, we developed a simple *in vivo* assay to directly demonstrate that HVEM/BTLA is an immune checkpoint for T-cell mediated tumor control.

**Conclusions:** Our results show that targeting HVEM is a promising strategy for cancer immunotherapy.

## Introduction

Immune escape by tumor is now considered a hallmark of cancer [1]. Many immune mechanisms are involved to explain the loss of tumor control, including defective MHC function and expression, recruitment of suppressive immune cells, and expression of co-inhibitory receptors such as PD-L1 [2]. In the last few years, targeting co-inhibitory molecules with antibodies has shown impressive results in tumor regression and overall survival, leading to the approval of anti-CTLA-4, anti-PD-1 and anti-PD-L1 in numerous cancers [3]. However, the success of immune checkpoint inhibitors (ICI) is still partial and many patients fail to respond. Limited tumor infiltrate (cold tumors) or low expression of the targeted molecule may explain the relative inefficiency of ICI [4,5]. To overcome these limitations, it is necessary to explore other pathways that might be involved in immune escape and that could complement actual therapies.

Recently, a new co-inhibitory pair has been highlighted in anti-tumor immune response: HVEM (Herpes Virus Entry Mediator, TNFRSF14) and BTLA (B and T lymphocyte attenuator) [6]. These two molecules can be expressed by many immune cells, including T-cells, in which signaling through BTLA is associated with inhibition of their activation [7,8]. Additionally, the HVEM network includes many additional partners, such as LIGHT, Herpes Simplex Virus-1 (HSV-1) glycoprotein D (gD), lymphotoxin α (LTα) or CD160 [6]. Like BTLA, binding of HVEM to CD160 on T-cells is associated with an inhibition of their activation [9]. In contrast, LIGHT is clearly a T-cell activator since transgenic expression of LIGHT in T cells leads to massive activation, especially in mucosal tissues [10]. On the other hand, stimulation of HVEM expressed by T-cells by any of its ligands is associated with proliferation, survival and production of inflammatory cytokines, such as IL-2 and IFN-γ [9,11]. Several clinical studies have shown that HVEM expression is upregulated in many types of cancers including colorectal cancers [12], melanomas [13], esophageal carcinomas [14], gastric cancers [15], hepatocarcinomas [16], breast cancers [17], lymphomas [18] or prostate cancer [19]. In these studies, high levels of HVEM expression by tumors were associated with a worse prognosis and lower survival. Moreover, HVEM expression by tumors was also associated with a reduction in the numbers of tumor-infiltrating leukocytes (TIL) [12,14,16].

Few studies have considered targeting the HVEM network to affect tumor growth. In fact, various strategies to inhibit HVEM expression or function lead to increased T cell proliferation and function in syngeneic tumor mouse models [14,20,21]. However, to our knowledge, no study to date has assessed the possibility to use a monoclonal antibody (mAb) to HVEM to favor the anti-tumor immune response in a humanized context *in vivo*. Herein, we investigated the therapeutic potential of a murine antibody targeting human HVEM in humanized mice grafted with various human tumor cell lines. To generate humanized mice, we used immuno-compromised NOD.SCID.γc^null^ (NSG) mice, which are deprived of murine T-, B- and NK- cells but that retain functionally immature macrophages and granulocytes [22]. We reconstituted these mice with human PBMC, allowing the effect of blocking HVEM to be studied on both tumors, murine myeloid cells and human T-cells.

## Methods

### Preparation of human peripheral mononuclear cells

Human peripheral blood mononuclear cells were isolated on a density gradient (Biocoll). Cells were washed in PBS 3% FCS and diluted at the appropriate concentration in 1X PBS before injection into mice.

### Humanized mice tumor model

All animals used were NSG mice (stock ≠005557) purchased from the Jackson Laboratory (USA). To assess therapeutic activity, 8–20-week-old NSG mice (males and females) were injected subcutaneously with 2.10^6^ tumor cells. One week later, mice were irradiated (2 Gy) and grafted the same day with 2.10^6^ huPBMC by retro orbital injection. Four to 5 days after transplantation, the anti-huHVEM antibody or isotype control was injected intra-peritoneally at 2 mg/kg. General state, body weight and survival of mice were monitored every 3-4 days to evaluate Graft-vs-Host-Disease (GVHD) progression. Mice were euthanized when exhibiting signs of GVHD, such as hunched back, ruffled fur, and reduced mobility. For CD8 depletion, mice were injected intra-peritoneally with 10mg/kg of the anti-CD8 MT807R1 (Rhesus recombinant IgG1 provided by the Nonhuman Primate Reagent Resource [23]) or the isotype control (clone DSPR1) the day following humanization, as previously described [24].

### Antibodies

The clone 18.10 has been described previously [25]. Briefly, 18.10 is a murine IgG1 anti-human HVEM mAb and was produced as ascites and purified by protein A binding and elution with the Affi-gel Protein A MAPS II Kit (Bio-rad). The mouse IgG1 isotype control (clone MOPC-21 clone), the rat IgG2b anti-Gr1 (clone RB6-8C5) and the isotype control rat IgG2b (clone LTF-2) were purchased from Bio X Cell (West Lebanon, NH, USA).

### Cell lines

PC3 (non-hormono-dependent human prostate cancer cells), Gerlach (human melanoma cells), MDA-MB-231 (breast cancer cells), DU145 (prostate cancer cells) were grown in high glucose DMEM media supplemented with 10% FCS, L-glutamine and antibiotics (Penicillin/Streptomycin). PC3 and MDA-MB-231 were genetically authenticated before the initiation of the experiments (Eurofins). All cells were confirmed to be free of mycoplasmas before injection into mice by the MycoAlert detection kit (Lonza). Tumor growth was monitored using an electronic caliper and volumes were determined using the following formula: [(length*width^2^)/2]. The PC3-GFP cell line was generated in the laboratory by lentiviral transduction (details available on request).

### Generation of HVEM deficient PC3 clone using CRISPR-Cas9 technology

50,000 PC3 cells were seeded in a 24-well plate. Twenty-four hours later, cells were incubated with sgRNA complementary to exon 3 of HVEM (GCCAUUGAGGUGGGCAAUGU + Scaffold, TrueGuide Synthtetic guide RNAs, Invitrogen™), Cas9 nuclease (TrueCut™ Cas9 Protein v2, Invitrogen™) and lipofectamine (Lipofectamine™ CRISPRMAX™ Cas9 Transfection Reagent, Invitrogen™) according to manufacturer instructions (TrueCut Cas9 protein v2 (27/09/2017)). After three days, efficiency was evaluated with GeneArt Genomic Cleavage Detection Kit (Invitrogen™) according to the manufacturer instructions. For this assay, DNA was amplified with the following primers: TGCGAAGTTCCCACTCTCTG (Forward) and GGATAAGGGTCAGTCGCCAA (Reverse). Cells were cloned by limiting dilution in 96-well plates. Clones were screened for HVEM expression by flow cytometry using anti-HVEM (clone 94801, BD) and were considered as negative if HVEM expression was undetectable for at least 3 subsequent measurements.

### In vitro assays

PC3 cells were seeded in 96-wells plate at 7000 cells/well in RPMI medium. Cells were treated by the anti HVEM antibody or its isotype control MOPC21 coated at 10μg/ml. Cell death was evaluated by flow cytometry after 16 hours of incubation (37°C, 5% CO2) by 7-AAD staining. Macrophages from NSG mice were obtained by peritoneal wash. The target to effector ratio was 1:5 for apoptosis monitoring. For live cell imaging, apoptosis of the PC3 GFP cell line was assessed using the annexin V red (cat n°4641, Sartorius). Culture was monitored every hour during 16 hours by Incucyte and overlapping of GFP (green) and apoptosis staining (red) was quantified and reported as number of apoptotic cells/well.

### Phenotypic analysis by flow cytometry

Tumors were digested with 0.84mg/mL of collagenase IV and 10μg/mL DNAse I (Sigma Aldrich) for 40min at 37°C with an intermediate flushing of the tissue. Cells were passed through a 100μm-cell strainer and suspended in PBS 3% FCS. To eliminate dead cells and debris, tumor cell suspensions were isolated on a Biocoll gradient. Rings were collected, washed, and cell pellets were suspended in PBS 3% FCS before counting on LUNA™ Automated Cell counter (Logos Biosystems). Subsequently, up to 2.10^6^ live cells were stained with viability dye (eF506, Fixable Viability Dye, ThermoFisher) for 12 min. at 4°C, Fc receptor were blocked with human FcR Blocking Reagent (120-000-442, Miltenyi Biotec) and anti-CD16/32 (clone 2.4G2) for 10 min. The followings antibodies were added for 35 min. at 4°C: hCD45-BUV805 (HI30, BD), hCD3-PECyn7 (SK7, BD), hCD4-PerCP (RPA-T4, Biolegend), hCD8-APC-H7 (SK1, BD), hKi67-AF700 (B56, BD), hCD270-BV421 (cw10, BD), and mCD45-BUV395 (30-F11, BD) hGranzymeB-APC (GB11, eBioscience), hPerforin-PE (B-D48, Biolegend) and mCD45-BUV395 (30-F11, BD). For intracellular staining, Foxp3/Transcription Factor Staining (eBioscience) or Cytofix/Cytoperm (BD) buffer sets were used. Cells were washed with 1X PBS before acquisition on an X20 cytometer (Becton Dickinson (BD), San Jose, CA). The absolute count of different populations was determined by adding 50 μL of Cell Counting Beads (Bangs Laboratories) before acquisition. Data were analyzed using FlowJo software (TreeStar, Ashland, OR, USA).

### NanoString nCounter expression assay

For Nanostring® experiment, 14 to 15 weeks-old NSG mice were humanized and treated with anti-HVEM or isotype. Day 28 post humanization, tumors were harvested and TIL were isolated as described above. To maximize mRNA recovery, TIL were pooled by treatment groups (4 mice in the anti-HVEM group and 5 in the isotype control group). Then, cells were stained with viability dye (eF506) and anti hCD45-APC (HI30, Biolegend). Live hCD45^+^ cells were sorted using Aria II cell sorter. After centrifugation, cells were suspended in RLT buffer (Qiagen®) before freezing at -80°C until analysis. Data were normalized through the use of NanoString’s intrinsic negative and positive controls according to the normalization approach of the nSolver analysis software (Nanostring). For the analysis, 287 genes with raw count higher than 55 and an absolute fold-change of at least 20% were retained. Enrichment analysis was performed with Metascape [26], the GSEA desktop application [27] and Ingenuity Pathway Analysis (IPA) (Qiagen). For Metascape analysis, genes up or down regulated were analyzed separately whereas all genes were included in the GSEA or IPA analyses. For GSEA analysis, enrichment was performed using the Hallmark v7.2 or the Canonical Pathways v7.2 gene sets from the Broad Institute. With that workflow, a False Discovery Rate (FDR) or a Family Wise Error Rate (FWER) less than 0.25 is deemed “significant”.

### Statistical analysis

All statistical tests were performed with Prism software (Graph Pad Inc, La Jolla, CA, USA). To compare ranks between two groups, the p-value was calculated with a non-parametric two-tailed Mann-Whitney t-test. Statistical modeling of tumor growth was performed by linear and non-linear regression using the exponential growth model. When necessary, the p-values of these tests are indicated on the figure panels. Statistical power of the analyses (alpha) was arbitrarily set at 0.05. No test was performed *a priori* to adequate the number of samples with statistical power.

## Results

### Targeting HVEM with a mAb improves tumor control of HVEM+ cell lines in humanized mice

We first determined whether HVEM could be targeted for therapy by the anti-HVEM 18.10 monoclonal antibody. For that, we implanted various tumor cell lines in NSG mice and grafted human PBMCs few days after. No differences in tumor growth were observed with mice grafted with the prostate cancer cell line DU145 or with the triple-negative breast cancer cell line MDA-MB-231, which did not express HVEM (Figure 1A-B). In contrast, a significant reduction of tumor growth was observed in mice grafted with the HVEM-positive patient-derived melanoma cell line Gerlach and the PC3 prostate cancer cell line (Figure 1C-D). To rule out that the effect of the mAb on tumor growth was due to other differences than HVEM expression, we generated an HVEM-deficient PC3 cell line (clone 1B11) using CRISPR-Cas9 ribonucleoprotein (RNP) transfection (Figure 1E). The treatment with the mAb was completely inefficient on the 1B11 cell line in humanized mice (Figure 1E). Of note, the knock-down of HVEM impacted tumor growth to the same extent than the mAb on HVEM^+^ PC3 cells (Figure 1D-E). Thus, HVEM expression on the tumor was mandatory for the therapeutic efficacy of the mAb.

**Figure 1:**
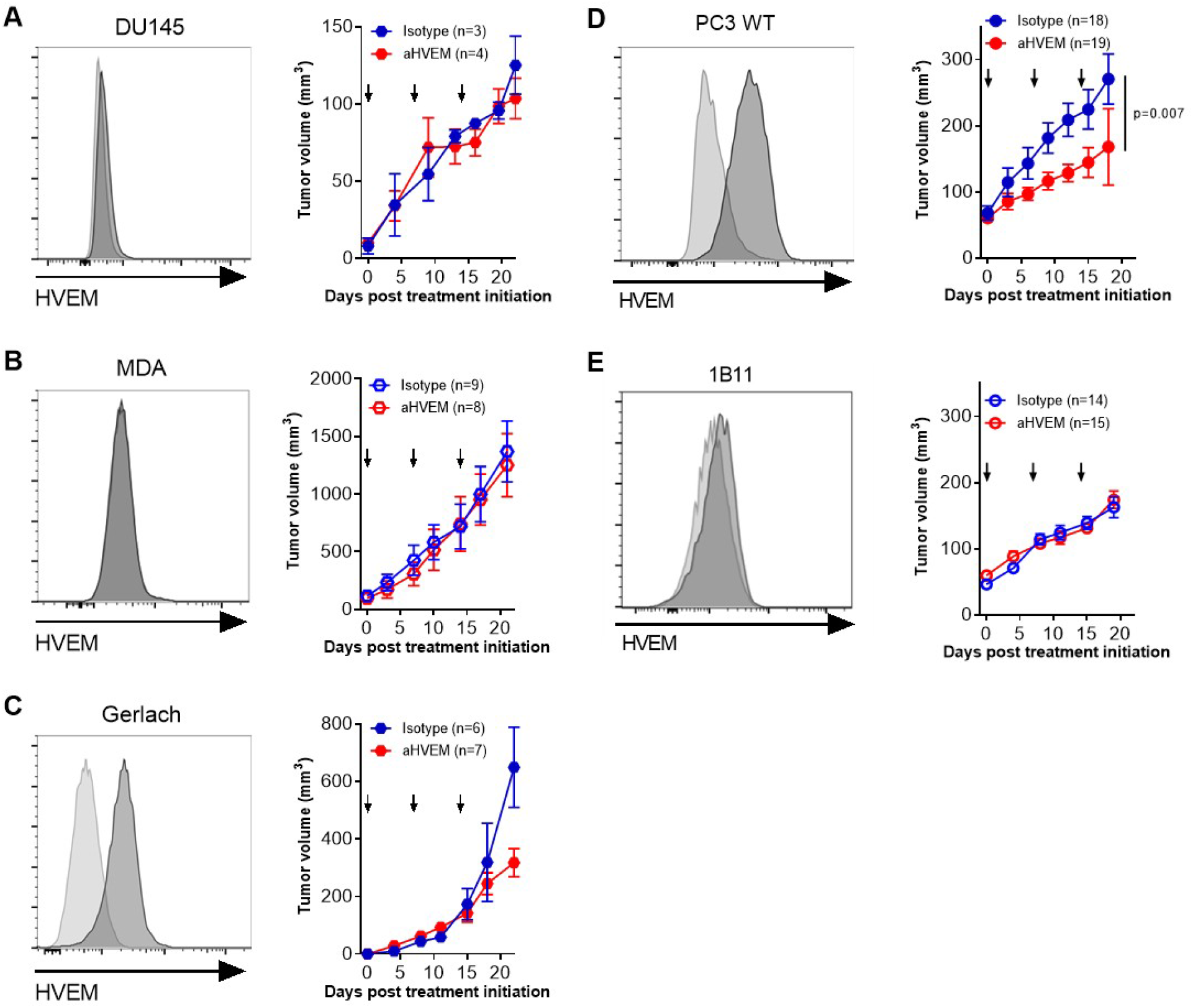
Targeting HVEM with a mAb improves tumor control of HVEM+ cell lines in humanized mice. HVEM expression and tumor growth of the prostate cancer cell line DU145 (A), the breast cancer cell line MDA-MB-231 (MDA) (B), the patient derived melanoma Gerlach (C), the prostate cancer cell line PC3 (D) and the HVEM-deficient PC3 clone 1B11. HVEM expression was determined by flow cytometry with the anti-HVEM mAb (clone 18.10) and a secondary antibody. Curves represent the mean tumor volume (±SEM) from one experiment with DU145 and at least two for the others. Numbers of mice at the beginning of the experiments are indicated in brackets. Arrows indicate the time of the injections. The p value on the graphs indicate the probability that the slopes are equal using a linear regression model.

### NSG myeloid cells are able to kill PC3 cells in presence of the anti-HVEM antibody

We next evaluated whether the mAb would mediate direct killing of tumor cells since HVEM has been linked to pro apoptotic signaling [28]. However, the anti-HVEM mAb was unable to induce tumor cell death *in vitro* (Figure 2A). In contrast, a significant reduction in tumor growth after mAb treatment was observed for the parental PC3 cell line in non-humanized NSG mice (Figure 2B). Because NSG mice are on a NOD genetic background which is deficient for complement activity [22], we surmised that innate immunity of NSG mice might be involved in the activity of the mAb. Indeed, depletion of monocytes and neutrophils with an anti-Gr1 mAb completely reverted the effect of the treatment, but a high mortality of NSG mice was observed (Figure S1). We thus co cultured PC3 cells with macrophages obtained from peritoneal lavage of NSG mice (Figure S2). Using live imaging, we observed a progressive increased proportion of apoptotic cells in presence of the anti-HVEM mAb (Figure 2C). Furthermore, video microscopy of the co-cultures revealed that tumor cells were killed by a cell-contact dependent mechanism with no evidence for engulfment of tumor cells (Video S1). Thus, innate immunity of NSG mice is not passive during treatment and may participate in tumor killing following treatment with the mAb.

**Figure 2:**
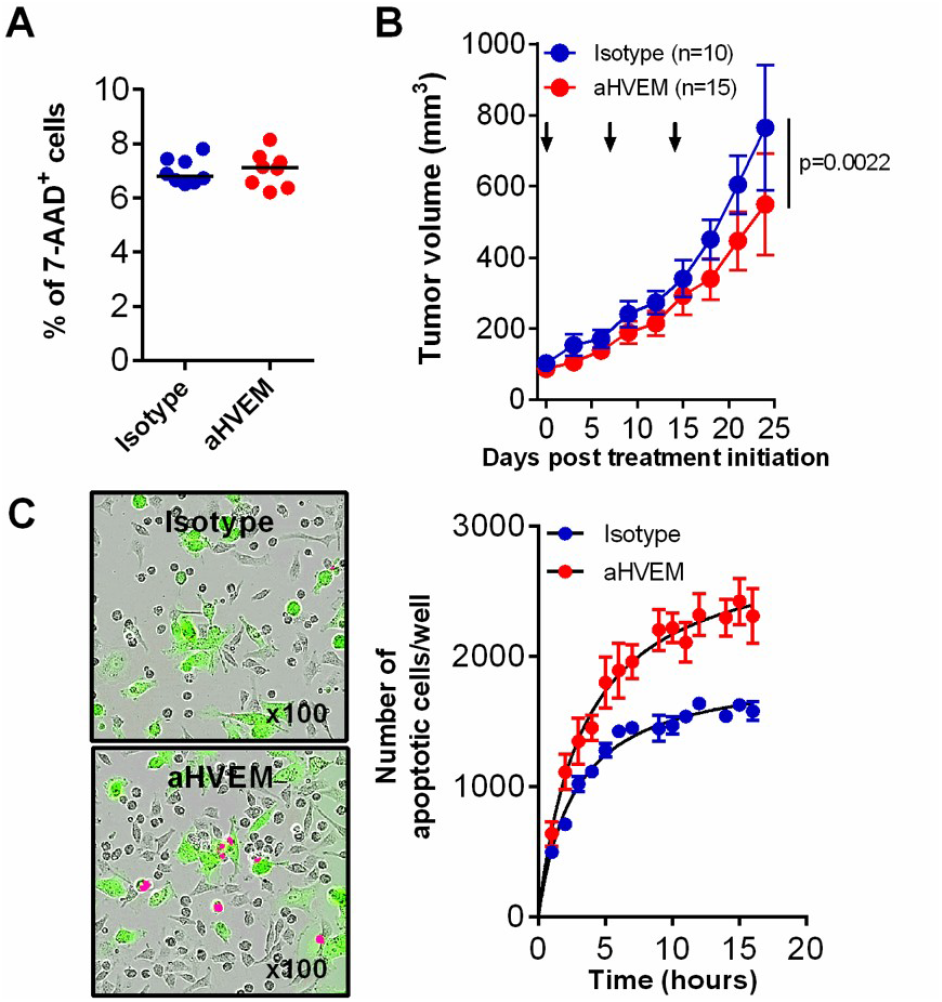
NSG myeloid cells are able to kill wild-type PC3 cells in presence of the anti-HVEM antibody. (A) Frequencies of 7AAD+ cells on the parental cell line PC3 in culture with anti-HVEM or isotype control mAb were determined by flow cytometry. (B) Tumor growth of the parental PC3 cell line in non-humanized NSG mice treated with the anti-HVEM or Isotype control mAb. Data are cumulative of 3 independent experiments. (C) GFP-expressing wild-type PC3 cells were co-cultured with NSG peritoneal macrophages and an apoptosis staining reagent. Magnification is indicated. (D) Overlap of GFP (green) and apoptosis staining (red) was quantified and reported as number of apoptotic cells/well ± SEM of technical replicates. Depicted are the results from one experiment.

### Treatment with the anti-HVEM mAb 18.10 results in an increase in TIL number and proliferation

To dig further into the mode of action of the mAb, we determined the relative frequencies of murine and human CD45^+^ cells in the tumor by flow cytometry. Among all CD45+ cells, murine cells were very rare compared to human cells. Human cells represented more than 90% of all CD45+ cells, in which CD3+CD4+ and CD8+ represented more than 95% (Figure 3A-C), showing that the tumor was mostly infiltrated by human T-cells. These proportions were not changed by the treatment. In contrast, we observed an increase in CD4 T-cells numbers and a similar tendency for CD8 T-cells in the anti-HVEM-treated group (Figure 3D). Additionally, frequencies of cells expressing the proliferation marker Ki67 were significantly elevated in both CD4 and CD8 T-cells (Figures 3E).

**Figure 3:**
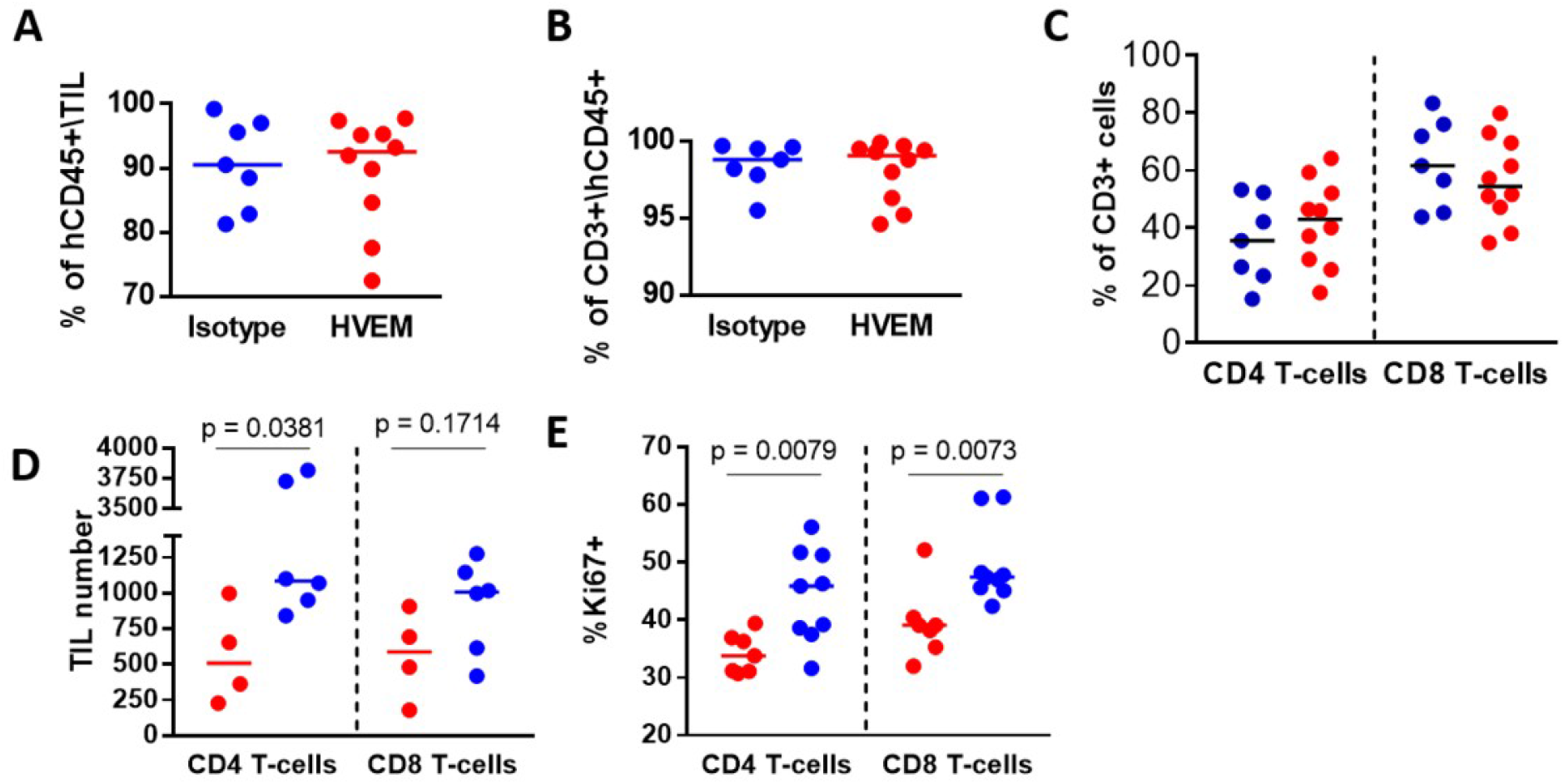
Treatment with the anti-HVEM mAb 18.10 results in an increase in TIL number and proliferation. Frequencies of human CD45+ cells among all CD45+ cells (A), of human CD3+ among human CD45+ cells (B) and of CD4 and CD8 T cells among human CD3+ cells (C) in the PC3 tumor were determined by flow cytometry. (D) Total number of CD4^+^ and CD8^+^ T-cells in PC3 tumors from one representative experiment out of 2. (E) Frequencies of Ki67-expressing cells among CD4^+^ and CD8^+^ T-cells in the tumor. Data are cumulative of two independent experiments performed at D21 post-humanization. Each dot is a mouse. The p values on the graphs indicate the probability that the median values were equals using the Mann-Whitney non parametric t-test.

### Tumor control is dependent on CD8^+^ T cells

To determine the contribution of CD8+ T cells to tumor control, we compared tumor growth in anti-HVEM-treated mice in mice depleted of their CD8+ T cells (Figure 4). Depletion of CD8 T-cells was clearly visible at the end of the experiment in the tumor (Figure 4A), showing that the depleting mAb had a long-lasting effect. Interestingly, depletion of CD8+ T cells before the initiation of the treatment reverted the effect of the anti-HVEM mAb on tumor growth (Figure 4B), showing that CD8 T cells were crucial for the therapeutic efficacy of the mAb. However, Granzyme B (GZMB) and Perforine 1 (PRF1) expression levels were not elevated in CD8^+^ T cells of the tumor of treated mice (Figure 4C-D), indicating that tumor control was dependent on CD8 T cell numbers rather than function.

**Figure 4:**
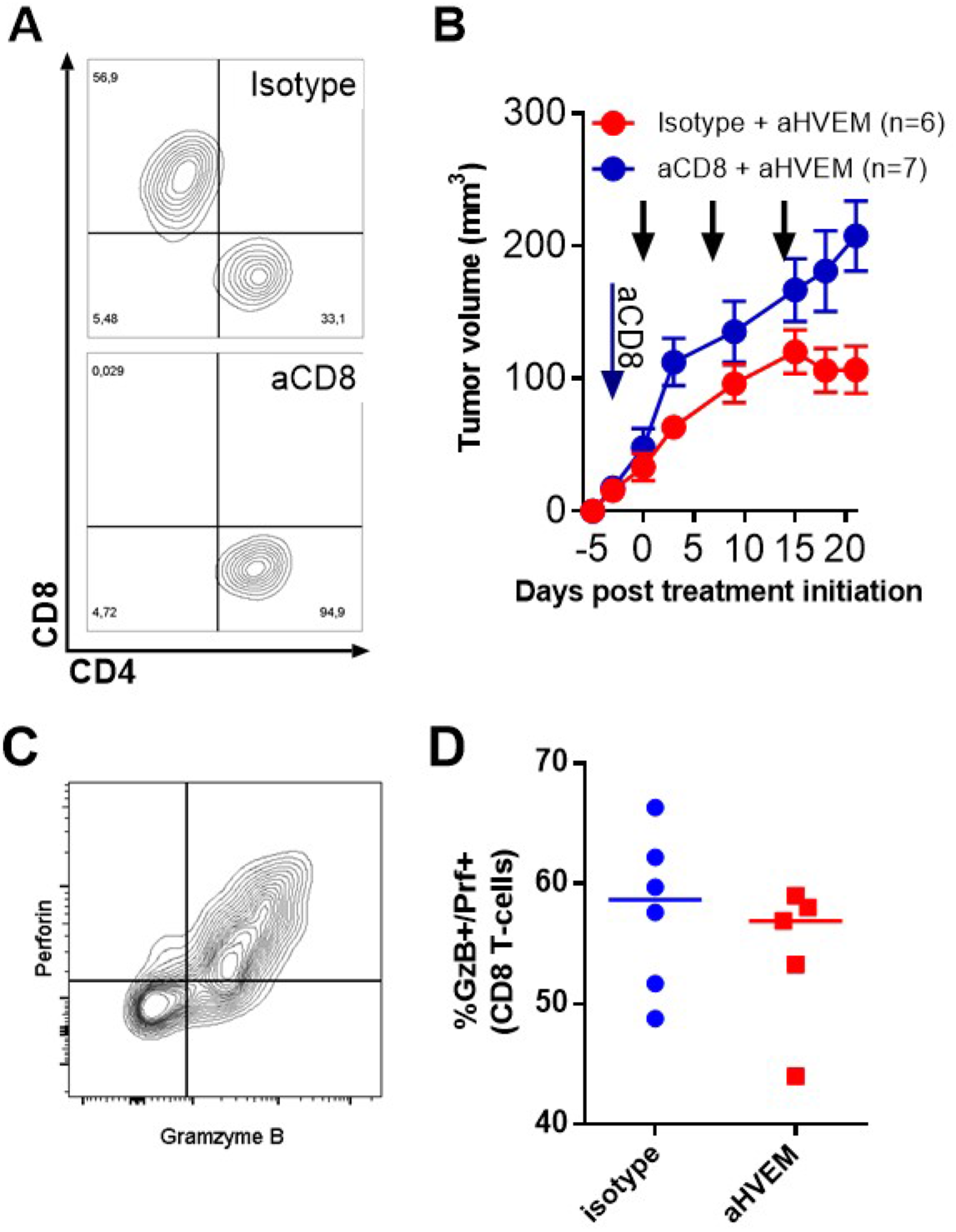
Tumor control by the mAb is dependent on CD8^+^ T cells. (A) Representative CD4/CD8 staining on human CD45+CD3+ T-cells in the tumor at the end of the experiment in a CD8-depleted (aCD8) or an isotype control treated mouse. (B) Growth of the PC3 cell line in humanized mice treated with anti-HVEM mAb and depleted or not of their CD8 T cells. CD8 T-cells were depleted on the day following humanization (blue arrow). Curves are the mean tumor volume (±SEM) in the indicated number of mice. Black arrows indicate the time of anti-HVEM mAb injection. Data are cumulative of two independent experiments.

### Treatment with the anti-HVEM mAb does not increase GVHD nor number or proliferation of human T cells

One possibility to explain these observations would be that the mAb behave as an agonist, directly activating human T cells *in vivo*, leading to better tumor control. Indeed, human T and B cells did express HVEM before injection into mice (Figure 5A). However, we observed similar weight loss and mortality in anti-HVEM or isotype control treated mice (Figure 5B-C), showing that GVHD induced by PBMC in NSG mice was not exacerbated by the treatment. Furthermore, the number and the proliferation status of human T cells in the spleens of treated animals were the same (Figure 5D-E). Our results show that anti-HVEM therapy in humanized mice reduced the growth of HVEM^+^ tumors by a mechanism that was independent of any agonist effect of the mAb.

**Figure 5:**
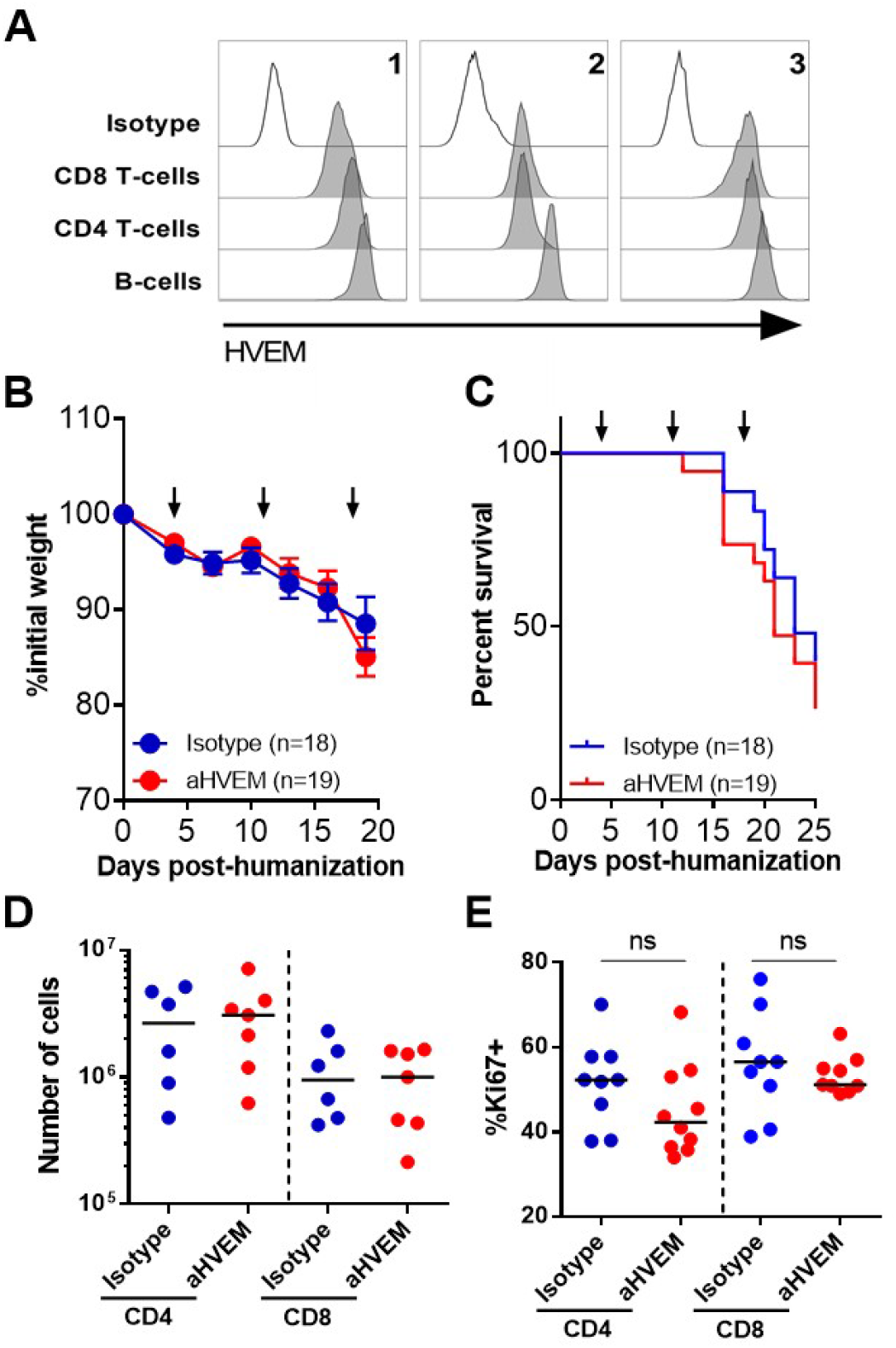
Treatment with the anti-HVEM mAb does not increase GVHD nor numbers or proliferation of human T cells. (A) HVEM expression in the indicated subsets was determined by flow cytometry on 3 different PBMC donors. Percentages of initial weight (B) and survival (C) of NSG mice following treatment by the anti-HVEM mAb or an isotype control are shown. Numbers (D) and frequencies of Ki67+ cells (E) in the indicated subsets.

### mRNA enrichment analysis showed increased activation and decreased immunosuppression in TIL of anti-HVEM treated mice

In order to better characterize the anti-tumor immune response following mAb treatment, we established a list of differentially expressed genes (DEG) in sorted hCD45^+^ TIL using the Nanostring Cancer Immune panel. Among the 730 genes included in the panel, 145 were up-regulated (log2FC>0.26) and 142 were down-regulated (log2FC<−0.3) in TIL from HVEM-treated mice (Figure S3). Of note, GZMB and PRF1 were among the genes with the highest levels of expression but the difference between the groups was weak, confirming our flow cytometry observation (Figure 4C-D). Moreover, several interleukins genes were enriched by the treatment, such as IL1A, IL7, IL22, EBI3, CSF2, and LTA, a ligand of HVEM. Likewise, the chemokines genes CCL5, CCL4, CCL1, CCL20 and others were enriched by the treatment. Finally, several members of the TNF super family were also enriched, such as TNFSF14 (LIGHT), another ligand of HVEM (Figure S3). Accordingly, unsupervised enrichment analysis revealed that up-regulated genes of the anti-HVEM group were enriched in members of several signatures related to interleukins/cytokines, and to activation signaling pathways, including the JAK-STAT, TNF-dependent NFkB and MAPK cascades (Figure 6A). Accordingly, NFKB1 and RELA were putative regulators of many genes of the DEG signature (Figure S4A). In addition, Gene Set Enrichment Analysis (GSEA) identified the “JAK-STAT signaling pathway” signature as significantly and positively enriched in TIL of HVEM-treated mice (Figure 6B).

**Figure 6:**
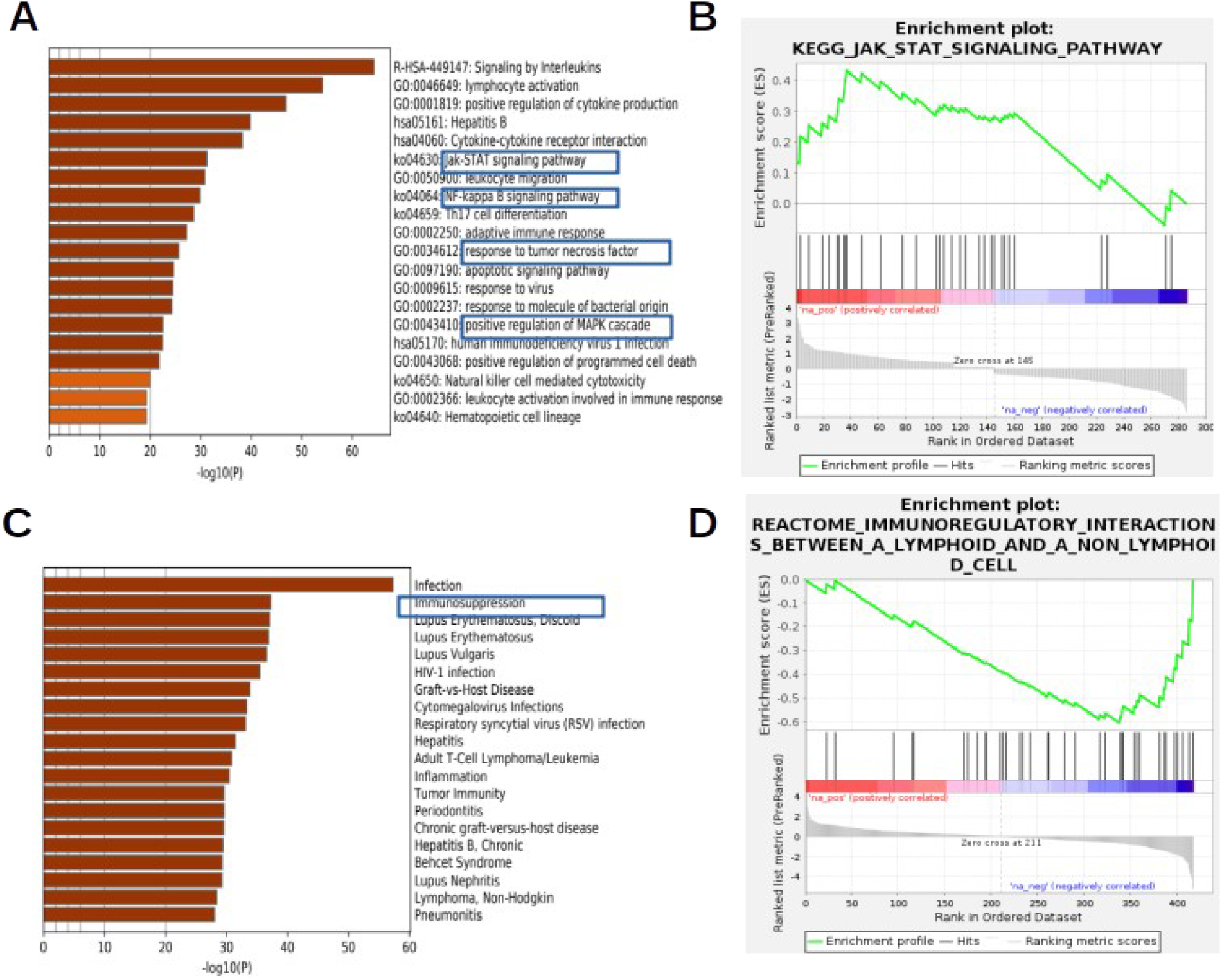
mRNA enrichment analysis show increased activation and decreased immunosuppression in TIL of anti-HVEM treated mice. Quantification of mRNA in CD45+ TIL of anti-HVEM or isotype-treated mice was performed with the Cancer Immune Nanostring panel in PC3-bearing humanized mice. (A) The first 20 terms significantly enriched in up regulated genes of CD45+ TIL of HVEM-treated mice are shown. (B) GSEA of the up-regulated genes identified the “JAK-STAT signaling pathway” signature as significantly enriched (p.val=0,01, q.val=0,26 (FDR), p.val=0,35 (FWER) in CD45+ TIL. (C) The first 20 terms significantly enriched in the genes down regulated by the anti-HVEM treatment are shown according to the DisGeNET database. (D) The “Immunoregulatory interaction between a lymphoid and a non lymphoid cell” signature was significantly enriched (p.val=0,002, q.val=0,148 (FDR), p.val=0,265 (FWER) in genes down modulated by the treatment.

On the other hand, some genes belonging to immuno-suppressive pathways were clearly down-regulated in HVEM-treated TIL such as ENTPD1 (CD39), IL10 and the co-inhibitory receptors BTLA, TIGIT, LAG3 and HAVCR2 (TIM3), as well as the “don’t eat me” receptor CD47 (Figure S3). Other cytokines/chemokines were also negatively affected by the treatment, such as IL21, IL13, CXCL13, TNFSF10 (TRAIL) or TNFSF113B (BAFF). Enrichment analysis of the genes down regulated in the anti-HVEM group using the DisGeNET database showed that the “Immunosuppression” signature was highly enriched in this gene set (Figure 6C). In addition, GSEA showed that genes belonging to the “immunoregulatory interactions between a lymphoid and a non lymphoid cell” signature was significantly depressed in the DEG signature (Figure 6D). In addition, IPA identified several “adhesion and/or binding of lymphocytes/leukocytes” signatures dependent on CSF2 and IL4 as the most significant biological functions associated with the DEG signature (Figure S4B). Overall, anti-HVEM treatment was associated with profound modifications of TIL, with an increased expression of genes belonging to activation and proliferation signaling pathways and a decreased expression of genes signing an exhausted phenotype.

### HVEM is an immune checkpoint during anti-tumor T cell immune response in humanized mice

To formally demonstrate that HVEM expression by the tumor was indeed an immune checkpoint, we devised a simple *in vivo* assay. We implanted the HVEM-positive or the HVEM-negative PC3 cells in NSG mice, and compared tumor growth with or without human PBMCs (Figure 7). Both cell lines grew equally well in non-humanized NSG mice (Figure 7A), showing that HVEM-deficiency did not impacted *in vivo* tumor development *per se*. In contrast, tumor growth of the 1B11 clone was reduced compared to the parental PC3 cell line in humanized mice (Figure 7B), directly showing that the lack of HVEM improved tumor control by human T-cells.

**Figure 7:**
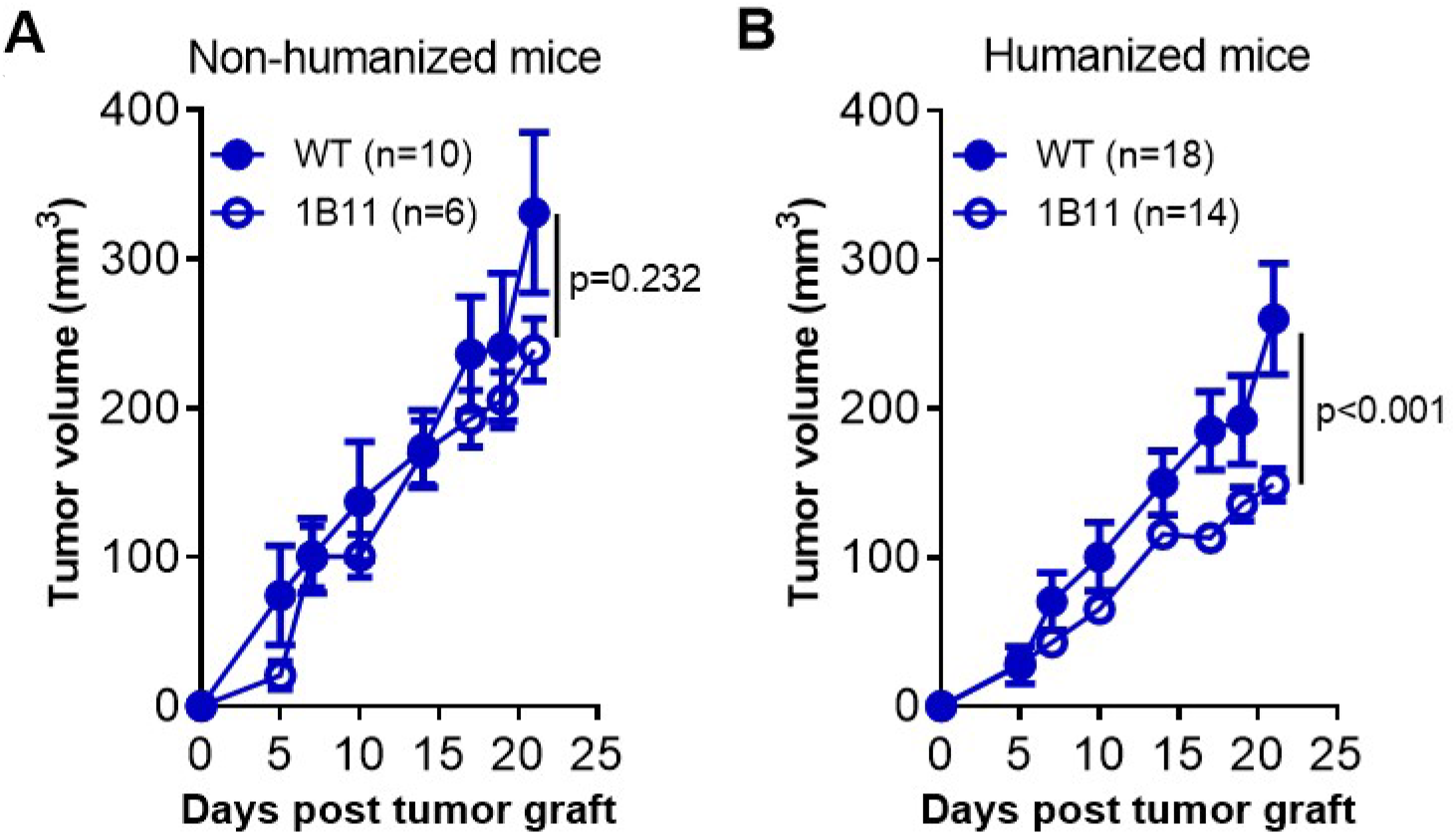
HVEM is an immune checkpoint during anti-tumor T cell immune response in humanized mice. Growth of the indicated PC3 cell lines (WT or 1B11) in non-humanized (A) or PBMC-humanized mice (B). Curves are the mean tumor volume (±SEM) in the indicated number of mice. Data are cumulative of at least two experiments. The p value on the graphs indicate the probability that the slopes are equal using a linear regression model.

## Discussion

Here, we report for the first time that HVEM can be targeted by a mAb to improve tumor control by human T cells *in vivo*. Moreover, we deciphered the mode of action of the mAb *in vivo* using complementary technologies. Furthermore, we developed a simple *in vivo* assay for immune checkpoint discovery and validation. To our knowledge, this is the first report that combine CRISPR/Cas9-mediated deletion of putative checkpoints in the tumor with assessment of tumor growth in humanized mice. One limitation of the assay is that PBMC-humanized mice are mostly reconstituted with T cells, as shown herein, limiting the usefulness of the assay to T cell-specific immune checkpoints. Nevertheless, we believe that this simple *in vivo* assay will be of great help to investigate other candidates in more advanced models of humanized mice. We show that the HVEM/BTLA checkpoint could be exploited for therapy in humanized mice using a mAb to human HVEM. We found that HVEM expression by the tumor was necessary and sufficient to elicit tumor control by the mAb, since the mAb had no effect on HVEM-negative cell lines and no agonist activity on human T cells. Park *et al.* showed in a syngeneic mouse model that transfecting an agonist scFv anti-HVEM in tumor cells resulted in increased T-cell proliferation, as well as improved IFN-γ and IL-2 production and better tumor control [20]. Aside the species differences, the discrepancy with our results could be explained by the fact that T-cells are strongly activated in huPBMC mice [29]. The down regulation of HVEM expression upon activation [30] may have limited the binding of the anti-HVEM antibody on T-cells in our model. Thus, it remains possible that the mAb would behave differently in humans. On the other hand, BTLA is up regulated upon T-cell activation [31], increasing the susceptibility of T-cells to inhibition by HVEM^+^ tumor cells [12,14,16,32]. We observed quite the opposite in the tumor micro environment following treatment, with an increase in HVEM and a reduction of BTLA gene expression, with a concomitant increase in LTA and LIGHT, two other ligands for HVEM. It is important to note that the binding sites of LIGHT and BTLA differ on HVEM [33]. So, the anti-HVEM mAb might have limited inhibition of activated T-cells through blockade of HVEM binding with BTLA but not with the other ligands that are T-cell activators. An alternative possibility would be that LIGHT and LTA in their soluble forms inhibit the interaction of HVEM with BTLA [34]. As of today, reciprocal regulation of HVEM and BTLA has not been reported but our observation is reminiscent of earlier findings showing reciprocal regulation of HVEM by LIGHT [30].

Previous studies in mice also showed that inhibiting HVEM expression on the tumor or its interaction with its ligands has a positive effect on T cells. Injection of a plasmid encoding a soluble form of BTLA (to compete with endogenous BTLA for HVEM) was associated with an increase in inflammatory cytokines production by TIL and a decrease in anti-inflammatory cytokines at the RNA level [21]. In the same line, vaccination to a tumor-associated antigen was more efficient if HVEM interactions with its ligands were blocked by HSV-1 gD, allowing regression of large tumor mass [35]. Moreover, silencing HVEM expression in the tumor with siRNA was also associated with an increase in CD8 T cells and inflammatory cytokine production in a murine colon carcinoma model [14]. In addition, use of siRNA to HVEM on ovarian cancer *in vitro* promoted T-cells proliferation and TNF-α and IFN-γ production [36]. Numerous results from our study also support increased T cell activation in the absence of HVEM/BTLA signaling: TIL from mice treated with anti-HVEM expressed higher levels of JAK-STAT, NFkB and MAPK signaling pathways that are well known inducers of proliferation, differentiation, migration and apoptosis. However, increase in TIL absolute numbers might not be enough to allow tumor rejection. Comparison between TILs from mice treated with the anti-HVEM or isotype control mAb also highlighted decreased expression of many co-inhibitory receptors genes (BTLA, TIGIT, LAG3 and TIM3 [37,38]) or with immunosuppressive functions (CD39 and IL10), suggesting a lower exhaustion status. Overall, we propose that the treatment with the anti-HVEM mAb allows better control of tumor growth by increasing the number of cytotoxic CD8 T-cells with a less exhausted phenotype. Our analysis also suggests that this may primarily impacts adhesion and binding in the tumor.

We also identified the impact of myeloid cells of NSG mice during immunotherapy, an overlooked issue when using the model. Our results are in line with published observations relating the crucial role of myeloid cells in tumor control upon immune checkpoints inhibitors treatment in syngeneic mouse models [39–41]. Because of the murine nature of the mAb, binding to murine Fc-receptors present on myeloid cells of NSG might have propelled the therapeutic efficacy of the mAb. In our setting, we used IgG1, that is reported to bind to CD16 (FcgRIII) and CD32 (FcgRIIB), activating and inhibitory receptors, respectively [42]. However, NOD background has been associated with a strong decrease in FcgRIIB expression by macrophages [43]. Consequently, activating FcgRIII might be the main receptor involved in FcR-dependent activity of murine myeloid cells in NSG mice. Several possibilities exist to explain tumor killing by myeloid cells, through antibody-dependent cellular phagocytosis (ADCP), local secretion of cytokines or free radicals, expression of FasL and many others [44,45]. We did not see evidence for ADCP on the video microscopy collected during the course of this study, which rather indicated that cell killing was mediated by a cell-contact dependent mechanism, the nature of which remains to be determined. Overall, our data suggest the following model to explain the anti-tumor activity of our anti-HVEM antibody in NSG mice: binding of the mAb on HVEM expressed by the tumor would activate tumor immunogenic cell death by murine myeloid cells, which together with blockade of the HVEM inhibitory network, would limit exhaustion and enhance proliferation and retention/migration of cytotoxic human T-cells. The recent success of ICI for cancer immunotherapy (anti-CTLA-4, anti-PD-1/PD-L1) has confirmed the hypothesis that the immune system can control many cancers. In light of the promising results reported herein, anti-HVEM therapy might be combined with ICI and/or chemotherapy to further enhance anti-tumor immunity.

## Abbreviations

HVEM: Herpes Virus Entry Mediator
BTLA: B and T Lymphocyte Attenuator
TIL: Tumor-Infltrating Leukocytes
TNFSF: Tumor Necrosis Factor Superfamily
NSG: NOD.SCID.γc^null^
ADCP: Antibody Dependent Cellular Phagocytosis
ICI: Immune Checkpoint Inhibitors
RNP: ribonucleoproteins
DEG: Differentially Expressed Genes
GVHD: Graft-Vs-Host-Disease
IPA: Ingenuity Pathway Analysis
GSEA: Gene Set Enrichment Analysis

## Declarations

### Ethics approval and consent to participate

Human peripheral blood mononuclear cells were collected by Etablissement Français du Sang from healthy adult volunteers after informed consent in accordance with the Declaration of Helsinki. Mice were bred in our animal facility under specific pathogen-free conditions in accordance with current European legislation. All protocols were approved by the Ethics Committee for Animal Experimentation Charles Darwin (Ce5/2012/025).

### Consent for publication

All authors concur with the submission of the article in its present form

### Availability of data and materials

Reagents and datasets are available from the corresponding author on reasonable request.

### Authors’ contributions

SB and NA performed the experiments, analyzed the data and contributed to the writing of the manuscript, DO provided essential reagents and edited the manuscript, GM designed the study, analyzed the data and wrote the manuscript.

## Acknowledgments

The authors would like to thank Olivier Bregerie and Doriane Foret for taking care of our mice, Dr Pukar KC for technical help, Sylvaine Just-Landi for preparing the 18.10 mAb, Dr Armanda Casrouge and Claude Baillou for cell sorting, and Dr Benoit Salomon for critical reading of the manuscript.

## Supplementary Figures

**Figure S1:**
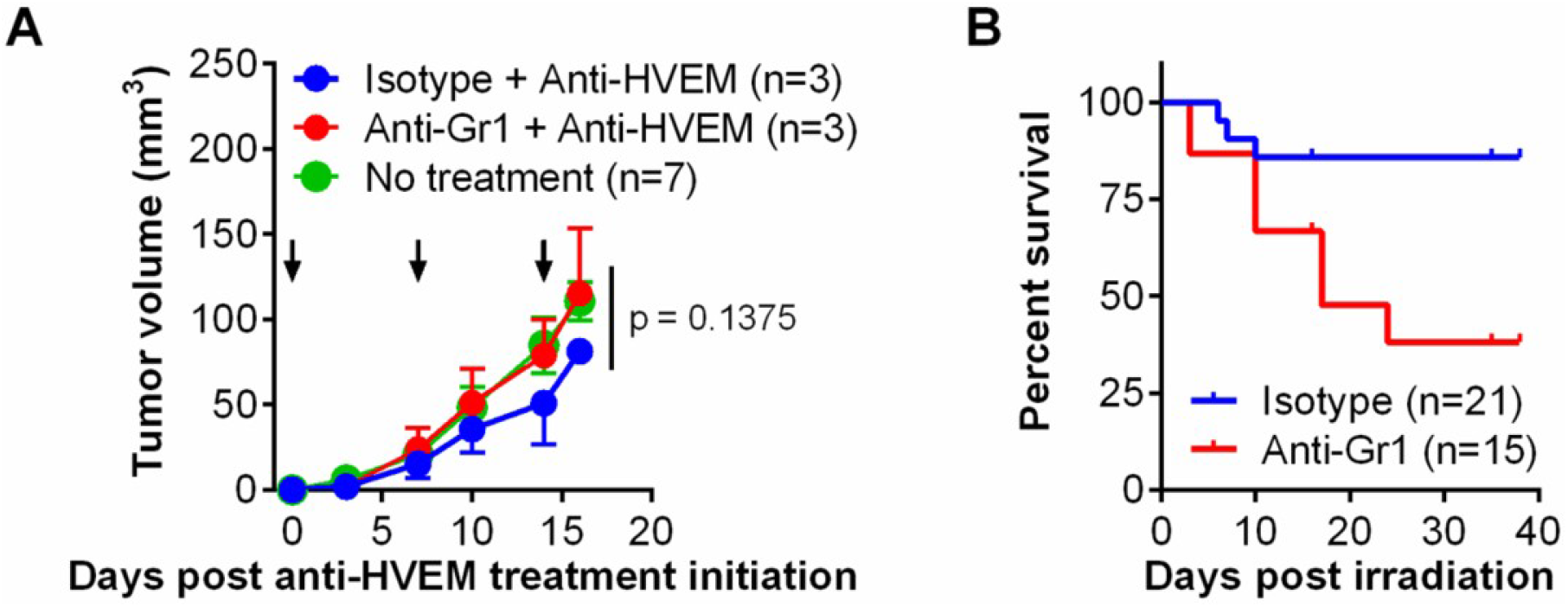
Depletion of Gr1+ cells reverted the effect of the anti-HVEM mAb in non humanized NSG mice. (A) Tumor growth of wild-type PC3 grafted in NSG mice treated with anti-HVEM (arrows) and anti-Gr1 (100μg) or isotype control twice a week. (B) Mice survival after treatment with anti-Gr1 mAb. Mice were treated with various doses of two different batches of anti-Gr1. Data are cumulative from 3 independent experiments.

**Figure S2:**
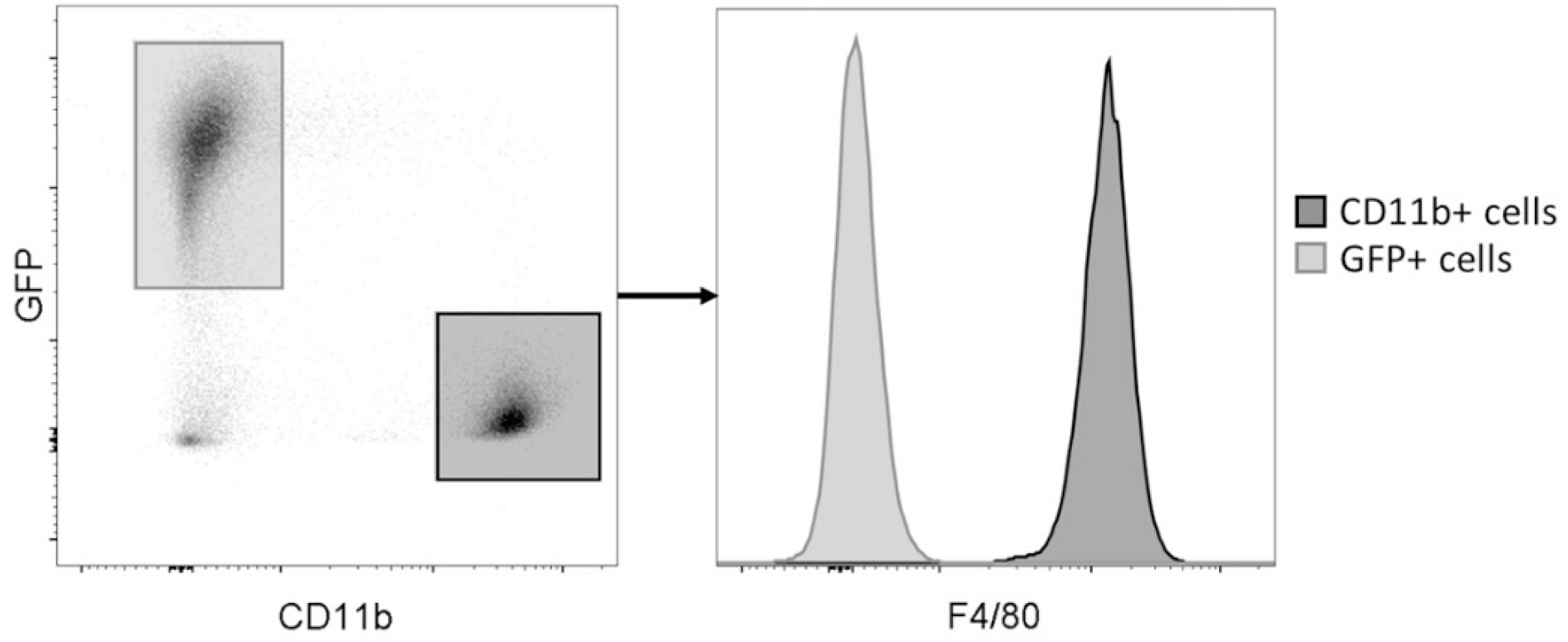
Co-cultures of GFP-PC3 cell line and cells from peritoneal lavage of NSG mice. Cells were stained with anti-CD11b and F4/80-specific mAb and analyzed by flow cytometry at the initiation of the co-culture.

**Figure S3:**
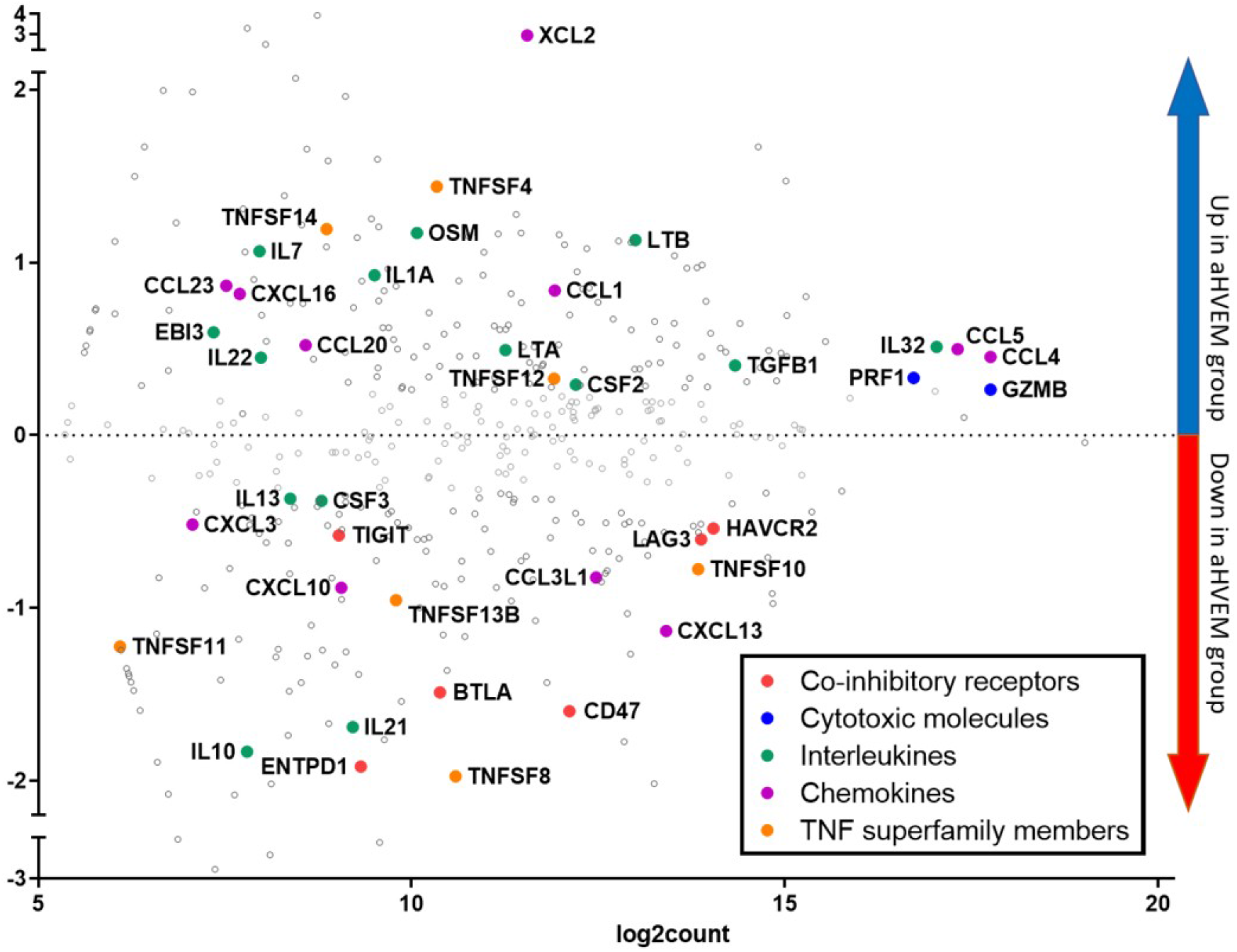
MA-plot comparing gene expression between TIL from aHVEM and isotype treated mice. Represented are the fold-change in the expression of a given gene between anti-HVEM- or isotype-treated mice (log2FC, y axis) vs the mean absolute count after normalization (log2count). Some notable genes are highlighted according to their biological functions by the indicated color code.

**Figure S4:**
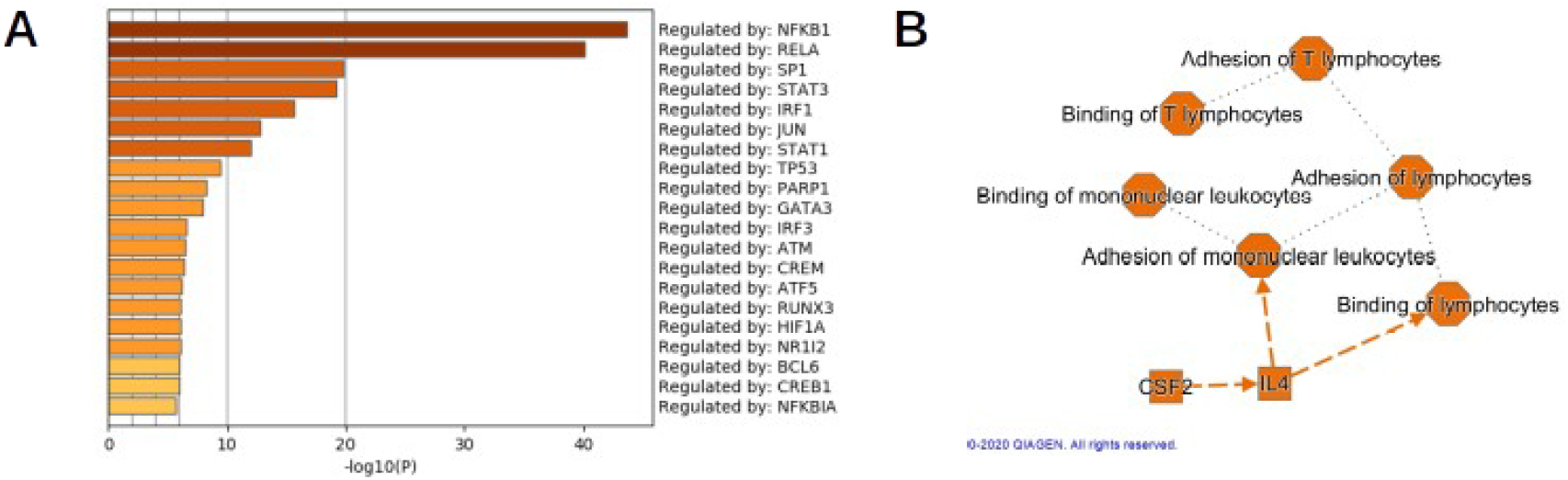
Enrichment analysis of DEG in TIL of anti-HVEM-treated mice. (A) Putative regulators of DEG were determined with Metascape and the TRRUST database (B) Representation of the most significant biological features from the DEG of anti-HVEM treated mice. The network was generated using the Graphical Summary algorithm of the Ingenuity Pathway Analysis software (Qiagen).

**Video S1: NSG macrophages are able to kill PC3 cell in presence of anti-HVEM by a cell-contact dependent mechanism.** GFP-expressing PC3 cells were incubating with NSG peritoneal macrophages and anti-HVEM or its isotype. Co-culture was followed by video microscopy overnight.

## References

1. Hanahan D, Weinberg RA. Hallmarks of cancer: The next generation. Cell. 2011;144:646–74.

2. Mittal D, Gubin MM, Schreiber RD, et al. New insights into cancer immunoediting and its three component phases-elimination, equilibrium and escape. Curr Opin Immunol. 2014;27:16–25.

3. Hargadon KM, Johnson CE, Williams CJ. Immune checkpoint blockade therapy for cancer: An overview of FDA-approved immune checkpoint inhibitors. Int Immunopharmacol. 2018;62:29–39.

4. Havel JJ, Chowell D, Chan TA. The evolving landscape of biomarkers for checkpoint inhibitor immunotherapy. Nat Rev Cancer. 2019;19:133.

5. Park Y-J, Kuen D-S, Chung Y. Future prospects of immune checkpoint blockade in cancer: from response prediction to overcoming resistance. Exp Mol Med. 2018;50:109.

6. Pasero C, Speiser DE, Derré L, et al. The HVEM network: new directions in targeting novel costimulatory/co-inhibitory molecules for cancer therapy. Curr Opin Pharmacol. 2012;12:478–85.

7. Cai G, Anumanthan A, Brown J a, et al. CD160 inhibits activation of human CD4+ T cells through interaction with herpesvirus entry mediator. Nat Immunol. 2008;9:176–85.

8. Sedy JR, Gavrieli M, Potter KG, et al. B and T lymphocyte attenuator regulates T cell activation through interaction with herpesvirus entry mediator. Nat Immunol. 2005;6:90–8.

9. Cheung TC, Steinberg MW, Oborne LM, et al. Unconventional ligand activation of herpesvirus entry mediator signals cell survival. Proc Natl Acad Sci U S A. 2009;106:6244–9.

10. Shaikh RB, Santee S, Granger SW, et al. Constitutive Expression of LIGHT on T Cells Leads to Lymphocyte Activation, Inflammation, and Tissue Destruction. J Immunol. 2001;167:6330–7.

11. Harrop JA, McDonnell PC, Brigham-Burke M, et al. Herpesvirus entry mediator ligand (HVEM-L), a novel ligand for HVEM/TR2, stimulates proliferation of T cells and inhibits HT29 cell growth. J Biol Chem. 1998;273:27548–56.

12. Inoue T, Sho M, Yasuda S, et al. HVEM Expression Contributes to Tumor Progression and Prognosis in Human Colorectal Cancer. Anticancer Res. 2015;35:1361–7.

13. Malissen N, Macagno N, Granjeaud S, et al. HVEM has a broader expression than PD-L1 and constitutes a negative prognostic marker and potential treatment target for melanoma. Oncoimmunology. 2019;8.

14. Migita K, Sho M, Shimada K, et al. Significant involvement of herpesvirus entry mediator in human esophageal squamous cell carcinoma. Cancer. 2014;120:808–17.

15. Lan X, Li S, Gao H, et al. Increased BTLA and HVEM in gastric cancer are associated with progression and poor prognosis. Onco Targets Ther. 2017;10:919–26.

16. Hokuto D, Sho M, Yamato I, et al. Clinical impact of herpesvirus entry mediator expression in human hepatocellular carcinoma. Eur J Cancer. 2015;51:157–65.

17. Tsang JYS, Chan K-W, Ni Y-B, et al. Expression and Clinical Significance of Herpes Virus Entry Mediator (HVEM) in Breast Cancer. Ann Surg Oncol. 2017;24:4042–50.

18. M’Hidi H, Thibult ML, Chetaille B, et al. High expression of the inhibitory receptor BTLA in T-follicular helper cells and in B-cell small lymphocytic lymphoma/chronic lymphocytic leukemia. Am J Clin Pathol. 2009;132:589–96.

19. Wang Q, Ye Y, Yu H, et al. Immune checkpoint-related serum proteins and genetic variants predict outcomes of localized prostate cancer, a cohort study. Cancer Immunol Immunother. 2020;

20. Park J-J, Anand S, Zhao Y, et al. Expression of anti-HVEM single-chain antibody on tumor cells induces tumor-specific immunity with long-term memory. Cancer Immunol Immunother. 2012;61:203–14.

21. Han L, Wang W, Fang Y, et al. Soluble B and T Lymphocyte Attenuator Possesses Antitumor Effects and Facilitates Heat Shock Protein 70 Vaccine-Triggered Antitumor Immunity against a Murine TC-1 Cervical Cancer Model In Vivo. J Immunol. 2009;183:7842–50.

22. Shultz LD, Schweitzer PA, Christianson SW, et al. Multiple defects in innate and adaptive immunologic function in NOD/LtSz-scid mice. J Immunol. 1995;154:180–91.

23. Schmitz JE, Simon MA, Kuroda MJ, et al. A nonhuman primate model for the selective elimination of CD8+ lymphocytes using a mouse-human chimeric monoclonal antibody. Am J Pathol. 1999/06/11. 1999;154:1923–32.

24. Petit NY, Lambert-Niclot S, Marcelin A-G, et al. HIV Replication Is Not Controlled by CD8+ T Cells during the Acute Phase of the Infection in Humanized Mice. PLoS One. 2015;10:e0138420.

25. Gertner-Dardenne J, Fauriat C, Orlanducci F, et al. The co-receptor BTLA negatively regulates human Vg9Vd2 T-cell proliferation: A potential way of immune escape for lymphoma cells. Blood. 2013;122:922–31.

26. Zhou Y, Zhou B, Pache L, et al. Metascape provides a biologist-oriented resource for the analysis of systems-level datasets. Nat Commun. 2019;10.

27. Subramanian A, Kuehn H, Gould J, et al. GSEA-P: A desktop application for gene set enrichment analysis. Bioinformatics. 2007;23:3251–3.

28. Pasero C, Barbarat B, Just-Landi S, et al. A role for HVEM, but not lymphotoxin-β receptor, in LIGHT-induced tumor cell death and chemokine production. Eur J Immunol. 2009;39:2502–14.

29. Ali N, Flutter B, Rodriguez RS, et al. Xenogeneic Graft-versus-Host-Disease in NOD-scid IL-2Rcnull Mice Display a T-Effector Memory Phenotype. PLoS One. 2012;7:10.

30. Morel Y, Schiano de Colella J-M, Harrop J, et al. Reciprocal Expression of the TNF Family Receptor Herpes Virus Entry Mediator and Its Ligand LIGHT on Activated T Cells: LIGHT Down-Regulates Its Own Receptor. J Immunol. 2000;165:4397–404.

31. Murphy KM, Nelson CA, Šedý JR. Balancing co-stimulation and inhibition with BTLA and HVEM. Nat Rev Immunol. 2006;6:671–81.

32. Sasaki Y, Hokuto D, Inoue T, et al. Significance of Herpesvirus Entry Mediator Expression in Human Colorectal Liver Metastasis. Ann Surg Oncol. 2019;26:3982–9.

33. Compaan DM, Gonzalez LC, Tom I, et al. Attenuating lymphocyte activity: The crystal structure of the BTLA-HVEM complex. J Biol Chem. 2005;280:39553–61.

34. Steinberg MW, Cheung TC, Ware CF. The signaling networks of the herpesvirus entry mediator (TNFRSF14) in immune regulation. Immunol Rev. 2011;244:169–87.

35. Lasaro MO, Sazanovich M, Giles-Davis W, et al. Active Immunotherapy Combined With Blockade of a Coinhibitory Pathway Achieves Regression of Large Tumor Masses in Cancer-prone Mice. Mol Ther. 2011;19:1727–36.

36. Zhang T, Ye L, Han L, et al. Knockdown of HVEM, a Lymphocyte Regulator Gene, in Ovarian Cancer Cells Increases Sensitivity to Activated T Cells. Oncol Res Featur Preclin Clin Cancer Ther. 2016;24:189–96.

37. Anderson AC, Joller N, Kuchroo VK. Lag-3, Tim-3, and TIGIT: Co-inhibitory Receptors with Specialized Functions in Immune Regulation. Immunity. 2016;44:989–1004.

38. De Sousa Linhares A, Leitner J, Grabmeier-Pfistershammer K, et al. Not All Immune Checkpoints Are Created Equal. Front Immunol. 2018;9.

39. Dhupkar P, Gordon N, Stewart J, et al. Anti-PD-1 therapy redirects macrophages from an M2 to an M1 phenotype inducing regression of OS lung metastases. Cancer Med. 2018;7:2654–64.

40. Gordon SR, Maute RL, Dulken BW, et al. PD-1 expression by tumour-associated macrophages inhibits phagocytosis and tumour immunity. Nature. 2017;545:495–9.

41. Gubin MM, Esaulova E, Ward JP, et al. High-Dimensional Analysis Delineates Myeloid and Lymphoid Compartment Remodeling during Successful Immune-Checkpoint Cancer Therapy. Cell. 2018;175:1014–1030.e19.

42. Bruhns P. Properties of mouse and human IgG receptors and their contribution to disease models. Blood. 2012;119:5640–9.

43. Luan JJ, Monteiro RC, Sautès C, et al. Defective Fc gamma RII gene expression in macrophages of NOD mice: genetic linkage with up-regulation of IgG1 and IgG2b in serum. J Immunol. 1996;157:4707–16.

44. Furness AJS, Vargas FA, Peggs KS, et al. Impact of tumour microenvironment and Fc receptors on the activity of immunomodulatory antibodies. Trends Immunol. 2014;35:290–8.

45. Gül N, Van Egmond M. Antibody-dependent phagocytosis of tumor cells by Macrophages: A Potent effector mechanism of monoclonal antibody therapy of cancer. Cancer Res. 2015;75:5008–13.

